# Inferring phage-bacteria infection networks from time series data

**DOI:** 10.1101/051581

**Authors:** Luis F. Jover, Justin Romberg, Joshua S. Weitz

## Abstract

In communities with bacterial viruses (phage) and bacteria, the phage-bacteria infection network establishes which virus types infects which host types. The structure of the infection network is a key element in understanding community dynamics. Yet, this infection network is often difficult to ascertain. Introduced over 60 years ago, the plaque assay remains the gold-standard for establishing who infects whom in a community. This culture-based approach does not scale to environmental samples with increased levels of phage and bacterial diversity, much of which is currently unculturable. Here, we propose an alternative method of inferring phage-bacteria infection networks. This method uses time series data of fluctuating population densities to estimate the complete interaction network without having to test each phage-bacteria pair individually. We use *in silico* experiments to analyze the factors affecting the quality of network reconstruction and find robust regimes where accurate reconstructions are possible. In addition, we present a multi-experiment approach where time series from different experiments are combined to improve estimates of the infection network and mitigate against the possibility of evolutionary changes to infection during the time-course of measurement.

## I. INTRODUCTION

Bacterial viruses are ubiquitous and play an important ecological role at the global scale. In the oceans, viruses are responsible for a significant fraction of bacterial mortality and as a result have an effect on global geobiochemical cycles [1–4]. By killing bacteria, they redirect resources from higher trophic levels and back into the microbial resource pool. Yet, not all bacteria types are susceptible to all virus types. Each phage type potentially infects subset of hosts which can be presented as complex networks of infection [5]. Quantifying who infects whom remains essential to understanding how individual-based traits affect ecosystem-wide properties in complex environments.

For more than 60 years, the host range of phage, i.e., the types of host that a phage type infects, has been measured using plaque assays [6]. A plaque assay is an experimental method in which a growing culture of bacteria on an agar surface are exposed to phage. Clear “plaques” are formed whenever the phage can infect and lyse the target host. Plaque assays are considered the gold-standard for determining infection but are hard to scale-up to community levels. The principal reason is that the majority of phage and bacteria in a community sample are not yet available in culture. In response, a number of (partially) culture-independent methods have been proposed, including viral tagging [7, 8], PhageFISH [9], and polonies [9]. Each of these methods requires some degree of culturing or co-visualization of labeled particles, which also presents challenges for scaling-up to complex communities. Moreover, none of these methods leverage the information contained in the temporal dynamics of virus-bacteria systems.

The inference of interaction networks from system dynamics is a field of study with wide-spread applications from inference of gene regulatory networks [10, 11], and chemical reaction [12], to neural networks [13]. The key insights from one class of inference methods is that statistical patterns in dynamics, including cross-correlation and mutual-information, can be leveraged to infer interaction [14]. However, such correlation-based approaches can be of limited value when applied to high dimensional systems with nonlinear interaction. As an alternative, Shandilya et al. [15] showed a method for reconstructing interaction networks from discrete measurements of the time series in systems where the underlying functional form of the interactions is known. Similarly, Stein et al. [16] following the work of Monier et al. [17] used discretized Lotka-Volterra equations to estimate interaction networks, model parameters, and time dependent perturbations in competitive microbial communities.

Here, we extend the approach of Stein et al. [16] to phage-bacteria systems with antagonistic interactions. We derive the principles underlying the method and test its validity using *in silico* experiments. As we show, inferring realistic phage-bacteria infection networks in complex communities may be possible given appropriate deployment of existing technologies already available to estimate changing genotype densities over time.

## II. METHOD

### A. Model

We model the interaction between *N*_*h*_ host types and *N*_*v*_ virus types using a generalization of the Lotka-Volterra predator-prey equations [18, 19]. The densities of multiple host and virus types are described by a system of differential equations that include the effect of competition between host types and the infection of host by multiple virus types [20, 21]:

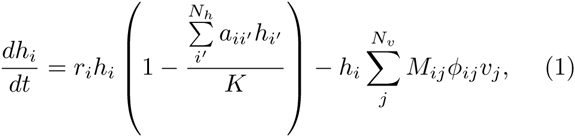

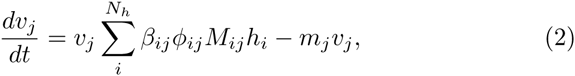

The model consists of *N*_*H*_ equations of the form (1) for the density of each host type, *h*_*i*_, and *N*_*v*_ equations of the form (2) for the virus densities, *v*_*j*_. In this system: *r*_*i*_ is the growth rate of host *i* in the absence of viruses and other hosts, *a*_*ii′*_ is the competitive effect of host *i′* on host *i*, *K* is the system-wide carrying capacity, *ϕ*_*ij*_ is the adsorption rate of virus *j* when attaching to host *i*, *β*_*ij*_ is the burst size of virus *j* when infecting host *i*, *m*_*j*_ is the decay rate of virus *j*. Finally *M*_*ij*_ is the infection matrix, i.e., a matrix representation of the infection network, which takes a value of 1 if host *i* is infected by virus *j* and zero otherwise.

### B. Numerical simulations of the dynamics; infection network ensembles and model parameters

To study the performance of our reconstruction method, we simulated time series of systems where several hosts and virus types interact. We used MATLAB’s ODE45 to numerically integrate systems of equations of the form described in Section II A. In doing so, we utilize both random infection networks and nested infection networks. Nested interaction networks are commonly observed in culture-based analyses, such that the host range of phage and the phage range of hosts form ordered subsets [22]. Following Jover et al. 2015 [23] we generated an ensemble of 100 infection matrices, each one with 10 host types and 10 virus types, spanning a spectrum of nestedness values. The infection matrices were generated by starting with a modular matrix and shifting interactions, through a random process, to regions that increase nestedness [23]. We also found feasible parameter sets (i.e., parameters with positive steady state densities) for each one of the infection matrices. We followed the procedure described in [23] to find feasible parameter sets. Namely, we select a subset of the model parameters and target densities (Table I) and use the steady state equations to solve for the rest of the parameters obtaining a feasible parameter set.

**TABLE I:** Parameter and target steady state density ranges used to find feasible parameter sets. Bacteria growth rates, *r*_*i*_, and virus decay rates, *m*_*j*_, were derived using the steady state equations and the parameters presented in this table (see Methods, given feasibility-based framework). The range denotes the limits of the uniform distributions used to generate parameters.

| Parameter (unit) | Range\Value |
| --- | --- |
| $\phi_j$ (ml/(virus · d)) | $10^{-8} - 10^{-7}$ |
| $\beta_j$ (viruses/cell) | 10 - 50 |
| $H_i^*$ (cell/ml) | $10^3 - 10^4$ |
| $V_j^*$ (virus/ml) | $10^6 - 10^7$ |
| $K$ (ml) | $\max(H_i^*) \times 100 = 10^6$ |

### C. Infection network reconstruction

Our method for reconstructing infection networks requires discrete measurements of the dynamics resulting from the interaction of different host and virus types. This method extends the approach described in [16] to host-phage systems. We will use only the equations describing the dynamics of the viruses (equations of the form (2)). We start by rewriting equation (2) in the form:

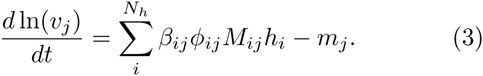

We assume that we have *N* + 1 measurements of the densities of all virus and host types in the system at times [*t*_1_, *t*_2_,…, *t*_*N*+1_]- For time step, Δ*t*_*n*_ = *t*_*n*+1_–*t*_*n*_, we can write a discretized form of equation (3):

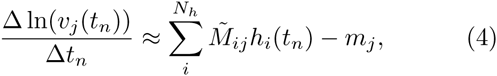

where we define the quantitative infection network 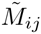:= *M*_*ij*_*ϕ*_*ij*_*β*_*ij*_,and 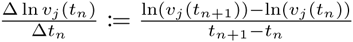. We can write an analogous equation to equation (4) for all time steps and all virus types in the system. All of these equations can be written in a compact form using a single matrix equation:

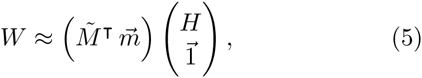

where *W* and *H* are matrices with elements *W*_*ij*_ = 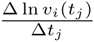 and *H*_*ij*_ = *h*_*i*_(*t*_*j*_), 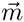 is the column vector of decay rates with elements *m*_*i*_, and 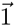 is a vector of ones with dimensions 1 × *N*. Given density measurements of the hosts and viruses we can reconstruct the quantitative infection network using equation (5). We solve the following minimization problem to obtain approximations 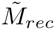 and 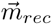 of the quantitative infection matrix, 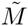, and the decay rate vector 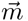:

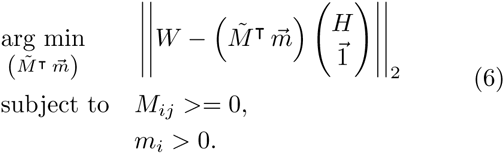

To solve this problem we used CVX, a package for specifying and solving convex problems [24, 25]. In this study we focus on the reconstruction of the quantitative infection network, but the method also infers decay rates for all virus types in the system. We use a normalized Frobenius distance between the original and reconstructed infection matrices as a metric of the quality of reconstruction, namely:

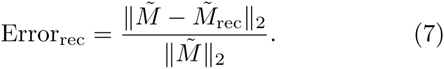

## III. RESULTS

### A. Reconstruction quality depends on the variability of the dynamics

We begin with an example in which there are 10 host types, 10 virus types and 20 virus-bacteria interactions. The effective infection rates (*ϕ* _*_ *β*) vary from 10^−7^ to 5 × 10^−6^. Figure 1 shows an example of a successful infection network reconstruction using the method described in Section II C. The matrices *W* and *H* were calculated using measurements of the dynamics every 6 min for a total of 96 hours. This results in a reconstruction error *Error*_*rec*_ = 0.01. The method is able to correctly identify all of the interactions. The small error arises from differences in the inferred quantitative values.

**FIG. 1:**
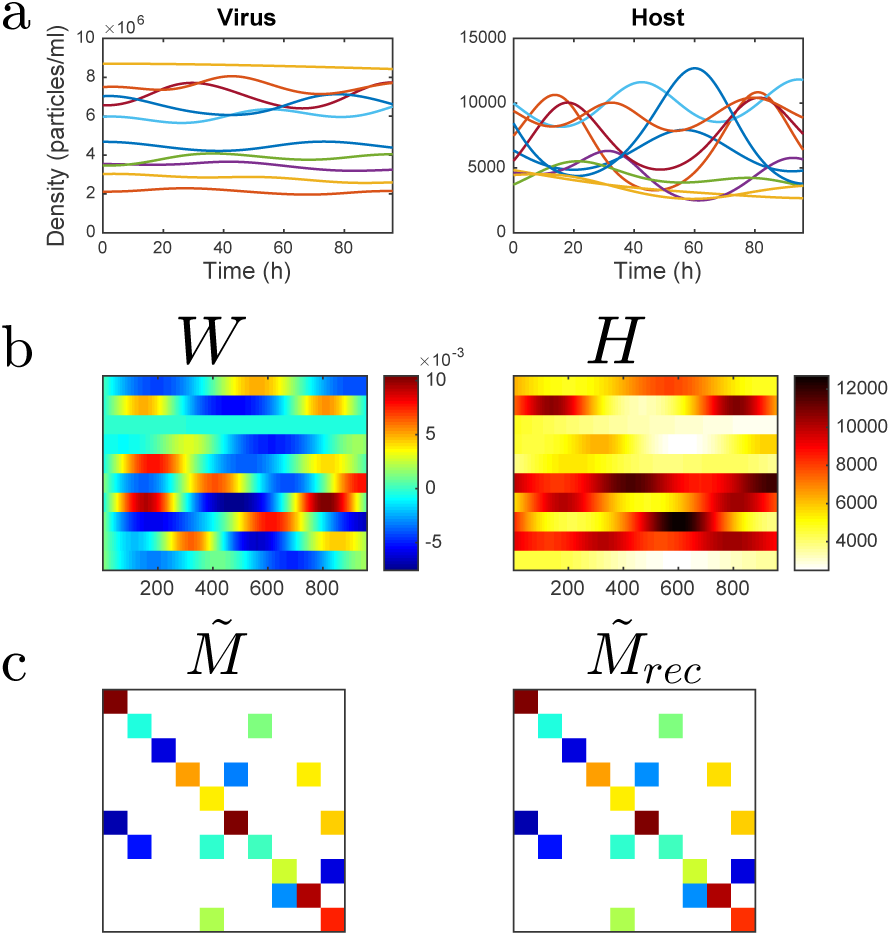
Example of infection network reconstruction, (a) Virus and host dynamics for 96 hours, (b) Matrices *W* and *H* constructed by taking measurements of virus and host densities every 6 min as described in Section II C. (c) Original and reconstructed infection matrices (*Error*_*rec*_ = 0.01). A feasible parameter set was used in the simulation as described in Section II B

In general, there are multiple factors affecting reconstruction quality. One important factor is the variability of the dynamics. For example, if the dynamics start at a fixed point, there would be no variability in the dynamics, the columns of the matrix *H* would all be identical and it would not be possible to infer the infection network. We test the effect of variability systematically by performing matrix reconstruction for an ensemble of matrices and different levels of variability. To control variability in the dynamics we change how far the initial densities are from the equilibrium densities. We initialize density of each host and virus type in the system at *x*_0_ = *x*_*eq*_(l ± *δ*), where *x*_*eq*_ is the equilibrium density of a given type and *δ* is a free parameter that controls the distance from its equilibrium density. We calculated the mean reconstruction error for an ensemble of 100 matrices (Figure 2). The reconstruction error has a maximum at *δ* = 0 (not shown for visualization purposes), which corresponds to starting the system at the equilibrium densities. The quality of the reconstruction increases as the initial conditions move away from the equilibrium densities.

**FIG. 2:**
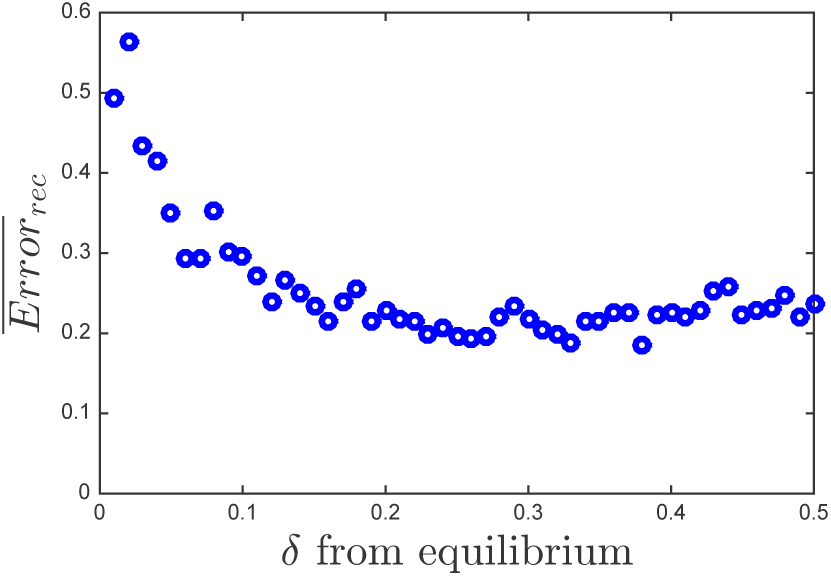
Mean reconstruction error as a function of the fraction away from the equilibrium densities, *δ*, for an ensemble of 100 matrices. Feasible parameter set were used in the simulation as described in section II B

### B. Reconstruction from multiple experiments: an alternative approach

We propose an improvement to the single experiment approach for reconstruction. In this alternative approach we combine measurements from different experiments to increase reconstruction quality. One key advantage of this approach is that, by increasing the number of experiments used for reconstruction, we can reduce the total time and number of measurements per experiment. This is a crucial advantage in virus-bacteria systems, which are known to evolve rapidly [26–28]. In the multiple-experiment approach we generate a host matrix *H* and a virus matrix *W* by combining matrices from multiple experiments that differ only in their initial conditions (Figure 3). This extends equation (5) to include information from multiple experiments. Specifically, assuming that we perform *p* different experiments and calculate matrices {*H*_1_, *H*_2_,…, *H*_*p*_} and {*W*_1_, *W*_2_,…, *W*_*p*_} for each experiment, we can write the system:

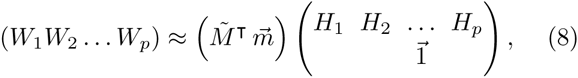

where 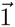 is a vector of ones with dimensions 1 × (*N*_1_ + *N*_2_+…+ *N*_*p*_), assuming that we take *N*_*i*_ measurements from experiment *i*. Using the same minimization process presented in Section II C we can obtain an approximation, 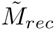, of 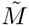.

**FIG. 3:**
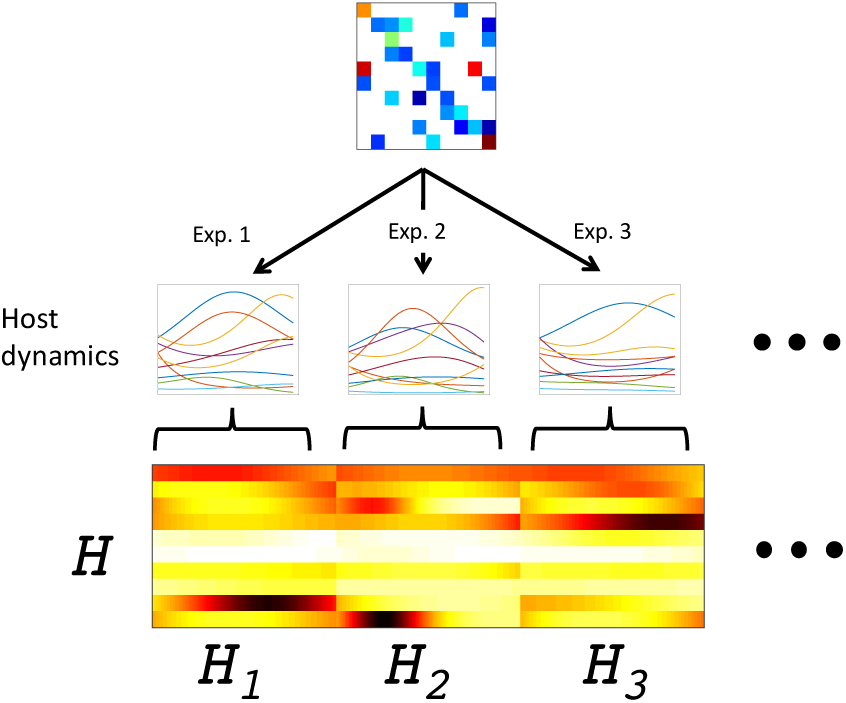
Schematic representation of how *H* is calculated in the multiple-experiment approach. Multiple experiments are performed with the same matrix 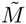 and different initial conditions.

Figure 4 compares the single and multiple experiments approach for three matrices with different nestedness values. We see how the multiple experiment approach results in lower reconstruction error for the three different cases. Figure 5 extends the comparison to an ensemble of 100 different matrices. We compare the multiple experiment approach to the average result of the single experiment approach. For a given matrix we performed 20 different experiments. Each experiment has the same infection matrix and the same model parameters but different initial conditions. We compare the performance of the reconstruction using each experiment individually vs. combining the measurements of the 20 experiments as described in equation (8). In this comparison we fix the total number of measurements; We compare the reconstruction error when using 960 measurements from a single experiment (measuring the dynamics every 6 minutes for 96 hours), against the performance when combining the first 48 measurement of all 20 experiments (every 6 minutes for 4.8 hours).

**FIG. 4:**
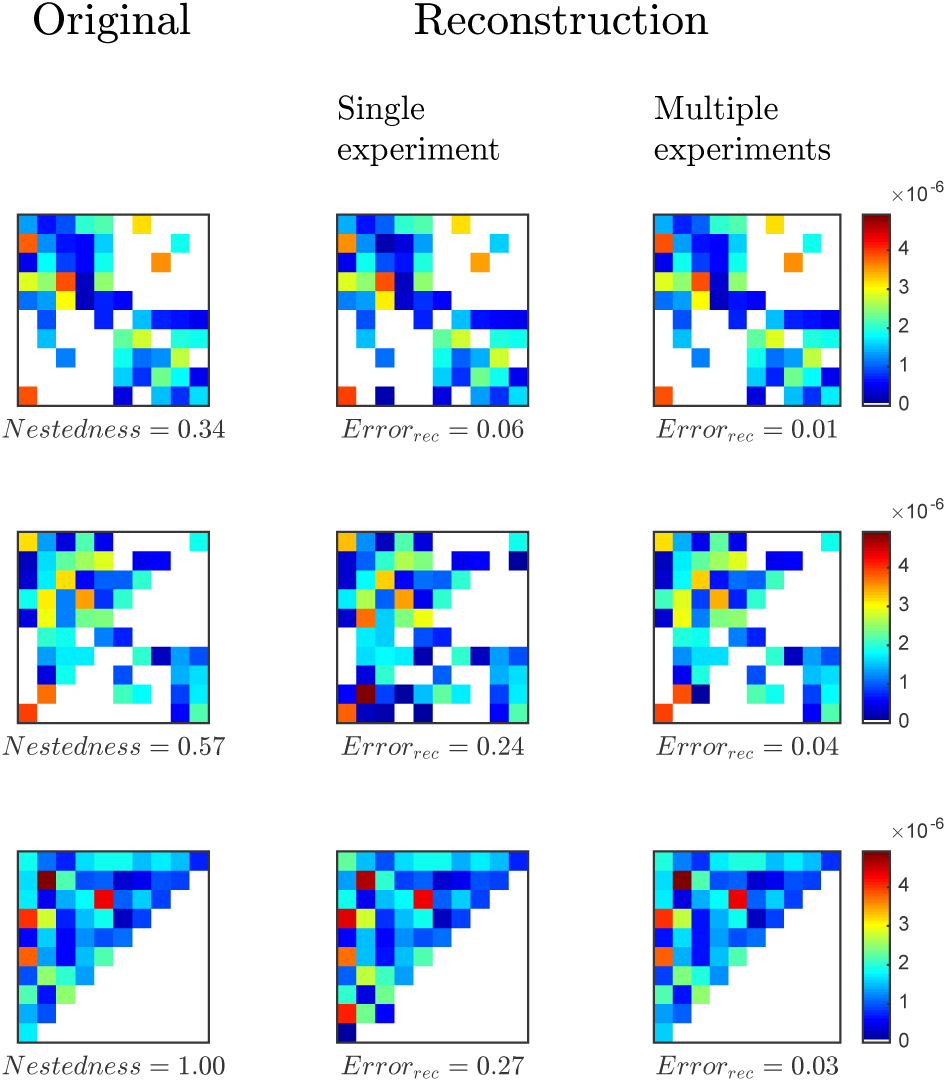
Examples of reconstruction for three different matrices and two different methods. Each row shows the original matrix and the resulting reconstruction for each method. The first column shows the original matrices with values of nest-edness (NODF): 0.34, 0.55, and 1 respectively. The middle column shows the reconstructed matrices and corresponding reconstruction errors for the single experiment approach using 960 measurements. The last column from the right shows the reconstructed matrices and corresponding errors for the multiple experiment approach using 20 experiments and 48 measurements per experiment. The total number of measurements is the same in the three different methods. The time between measurements is, Δ*t* = *6min*.

**FIG. 5:**
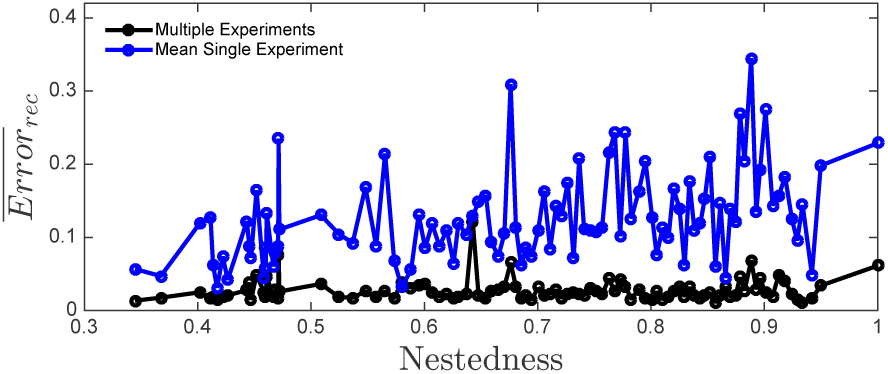
Reconstruction error vs Nestedness for two different methods. Black line denotes the reconstruction error, *Error*_*rec*_, using the multiple-experiments approach. Blue line describes the mean reconstruction error for the same 20 experiments used in the multiple-experiment approach but using each experiment separately. The total number of measurements is the same in both approaches.

We performed the comparison for 100 different matrices (Figure 5). Multiple-experiment reconstruction results in lower error than the average single experiment reconstructions across a wide range of nestedness values. The multiple experiment approach is also more robust; it results in smaller variance in the reconstruction error. Performing more than a few experiments not only decreases the mean reconstruction error, but also decreases the standard deviation significantly (Figure 6). For the specific configuration studied here reconstruction error minimizes around 18 experiments.

**FIG. 6:**
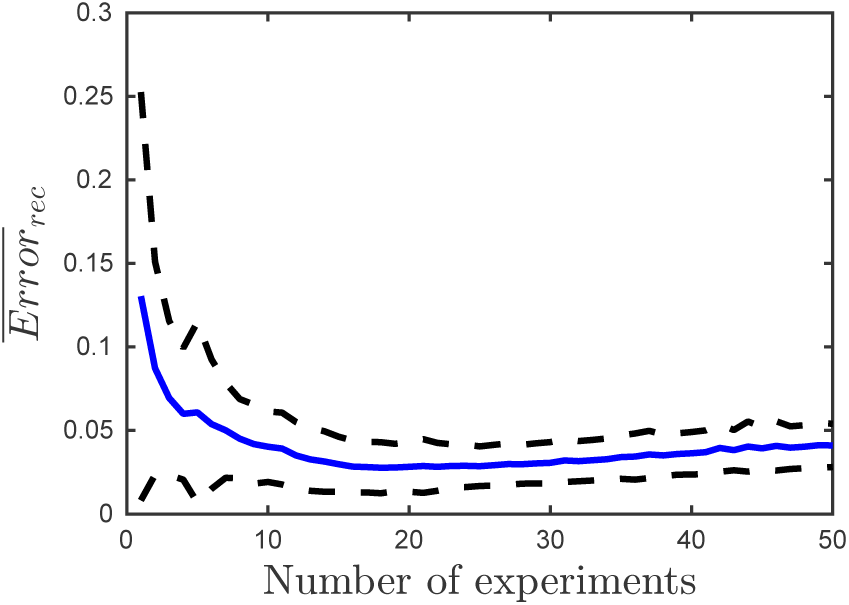
Mean (blue line) and standard deviation (dotted line) of the reconstruction error for 100 infection matrices as a function of the number of experiments used in the multiple-experiment approach. Fixed number of total measurements (960). Δ*t* = *6min*.

### C. Robustness of inference given noise in measurement

Here we evaluate the effect of measurement of white Gaussian noise on the quality of the inference. We follow the same procedure as in the noiseless case to reconstruct infection networks using multiple experiments. Figure 7 shows mean reconstruction error for an ensemble of 100 matrices as a function of the signal-to-noise ratio (SNR). We see that using 20 experiments and 48 measurements per experiment, network inference is possible for large signal-to-noise ratio, but reconstruction error increases significantly when the noise approaches 10% of the signal (SNR = 10 dB).

**FIG. 7:**
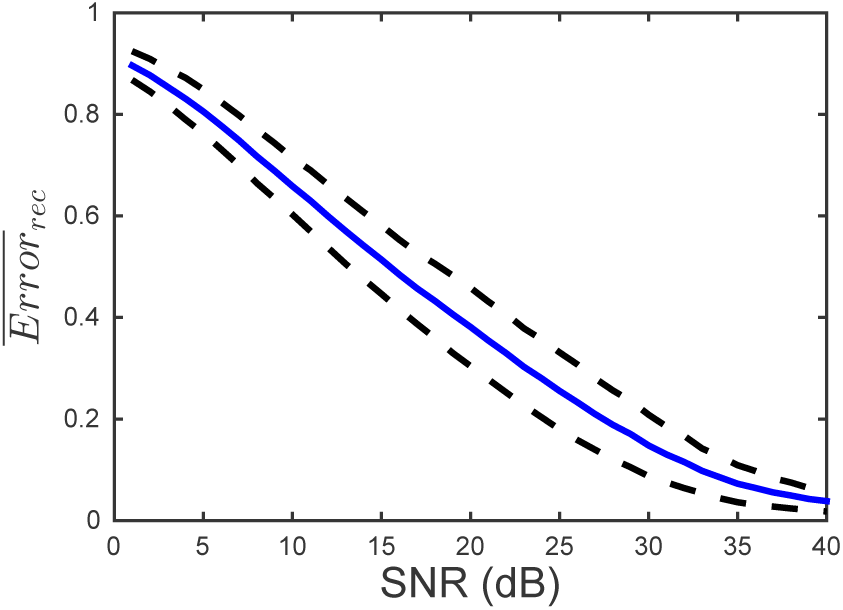
Mean (blue line) and standard deviation (dotted line) of the reconstruction error for 100 different matrices as a function of the signal-to-noise ratio. The multiple experiment approach was used to reconstruct the matrix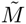. For each reconstruction, the matrices *H* and *W* were constructed using 20 runs with different initial conditions and 48 measurements per run. Δt = 6 min

## IV. DISCUSSION

We presented a theory-driven method to estimate host-phage infection networks in an community with multiple virus and host types. Current experimental techniques to measure such networks are difficult to scale to large systems. In addition, techniques that depend on isolation of viruses and/or hosts capture only a subset of potential interactions present in natural environments. Our approach addresses these limitations by using time-series measurements of experiments involving the whole virus-bacteria community. We also presented an improvement over the single experiment approach for infection network reconstruction. In the multiple-experiment approach we combined measurements from multiple experiments increasing the variability and lowering the reconstruction error. The multiple-experiment approach has the additional advantage of requiring shorter measurement time per experiment. As a consequence, there is a lower probability of a host gaining resistance to a virus type or a virus developing the ability to infect a new host, increasing the chances of reconstructing the infection network of the target community.

The current method takes as input the measured densities of bacteria and phage in an environmental sample. Next-generation high-throughput sequencing techniques provide a means to characterize bacterial and viral communities in a variety of environmental samples [29–33]. In the past, such characterization has focused on phylogenetics groups, by using RNA and other marker genes. Such markers are insufficiently resolved with respect to differences in relevant phenotypes, e.g., phage-bacteria infectivity. However, new computational approaches are increasingly able to resolve strain-level dynamics from metagenomic datasets [34, 35]. The increased used of quantitative piplines from sample to strain *density* for both bacteria and viruses will enable the kind of inference proposed here.

Our present approach uses the nonlinear dynamics of virus populations, to infer virus-bacteria infection networks. Nonetheless, this method can be expanded by including nonlinear bacterial population dynamics to infer competitive interactions between bacteria types and bacterial growth rates. In developing this method, it is important to keep in mind that the present approach is adapted to a specific functional form of the interactions in a virus-bacteria communities. Experimental verification (e.g., see Stein et a l. [16]) is necessary to test whether or not the dynamical model is a sufficiently robust representation of naturally occurring systems. Nevertheless, this study presents key steps towards an alternative way of determining who infects whom in a virus-bacteria community. This view has the potential to significantly reduce the experimental burden, e.g., we are able to infer *n*_*h*_ × *n*_*v*_ interactions by measuring the dynamics of *n*_*h*_ + *n*_*v*_ organisms, and to overcome the limitations of culture-based approaches by inferring interactions without culturing.

## V. ACKNOWLEDGMENTS

The authors thank Sam Brown and Joey Leung for their comments and suggestions, and acknowledge the support of the NSF grant No.OCE-1233760.

